# I-CONVEX: Fast and Accurate *de Novo* Transcriptome Recovery from Long Reads

**DOI:** 10.1101/2020.09.28.317594

**Authors:** Sina Baharlouei, Meisam Razaviyayn, Elizabeth Tseng, David Tse

## Abstract

Long-read sequencing technologies demonstrate high potential for *de novo* discovery of complex transcript isoforms, but high error rates pose a significant challenge. Existing error correction methods rely on clustering reads based on isoform-level alignment and cannot be efficiently scaled. We propose a new method, I-CONVEX, that performs fast, alignment-free isoform clustering with almost linear computational complexity, and leads to better consensus accuracy on simulated, synthetic, and real datasets.

---

Alternative splicing is the process by which a single gene can create different alternative spliced forms (isoforms) by using different combinations of exons. The process of identifying isoforms is called transcriptome sequencing. Transcriptome sequencing methods fall into two categories: genome-guided and *de novo*. Genome-guided methods align reads back to the reference genome to identify the exon boundaries. This alignment information is often combined with reference annotations to assemble the transcripts. *De novo* transcriptome sequencing, on the other hand, uses information from the reads alone and does not rely on a reference genome. The *de novo* approach is not biased by the reference genome/annotation and thus can be used in applications with the mutated genome, such as cancer, or when a high-quality reference genome is not available.

Most transcripts are 1-10 kb long, and different isoforms can share the same subset of exons. Thus, accurate characterization of the exon connectivities using short reads (100-250 bp) is computationally challenging and in some cases, even statistically impossible^1–4^. In contrast, the transcriptome sequencing problem through long reads is statistically identifiable (**Supplementary Note 1**). However, such a task is computationally challenging due to higher error rates of long reads. To deal with the high error rate, various transcriptome sequencing pipelines^5–7^ have been developed^5,8^ and used to discover novel isoforms^9^, cancer fusion genes^10^, and genotypes of immune genes^11^.

**Figure 1a** illustrates the process of full-length transcriptome sequencing. Each long read covers a transcript completely, with substitution, insertion, and deletion errors distributed randomly. The number of reads covering each transcript depends on its abundance in the sequencing library. We define the *de novo* transcriptome recovery as the problem of using the full-length reads with random errors to estimate the sequence of the transcripts and their abundances. One solution to this problem is to cluster the reads based on their similarity, assuming that each cluster contains reads coming from the same isoform and within-cluster differences solely come from sequencing errors. The software ICE^5^, which is based on this clustering viewpoint, performs pairwise alignment among the reads to construct a similarity graph and then uses this graph to cluster the reads. While optimal clustering algorithms often require solving mathematically non-convex and computationally intractable problems, heuristic clustering algorithms such as maximal decomposition can be used in practice^5^. Unfortunately, these heuristics often provide no statistical guarantee for the final clusters. In addition, computing similarity graphs relies on aligners which are subject to parameterization and sensitivity/specificity tradeoffs. Another software based on this clustering viewpoint is IsoCon^8^. IsoCon first creates a nearest neighbor graph based on the pairwise edit distance of the reads, then it successively removes and denoises nodes with the largest number of neighbors. This procedure is continued until all the reads are clustered. While IsoCon demonstrates significantly better recall and precision compared to ICE, it is not scalable to large-scale datasets with millions of long reads.

**Figure 1:**
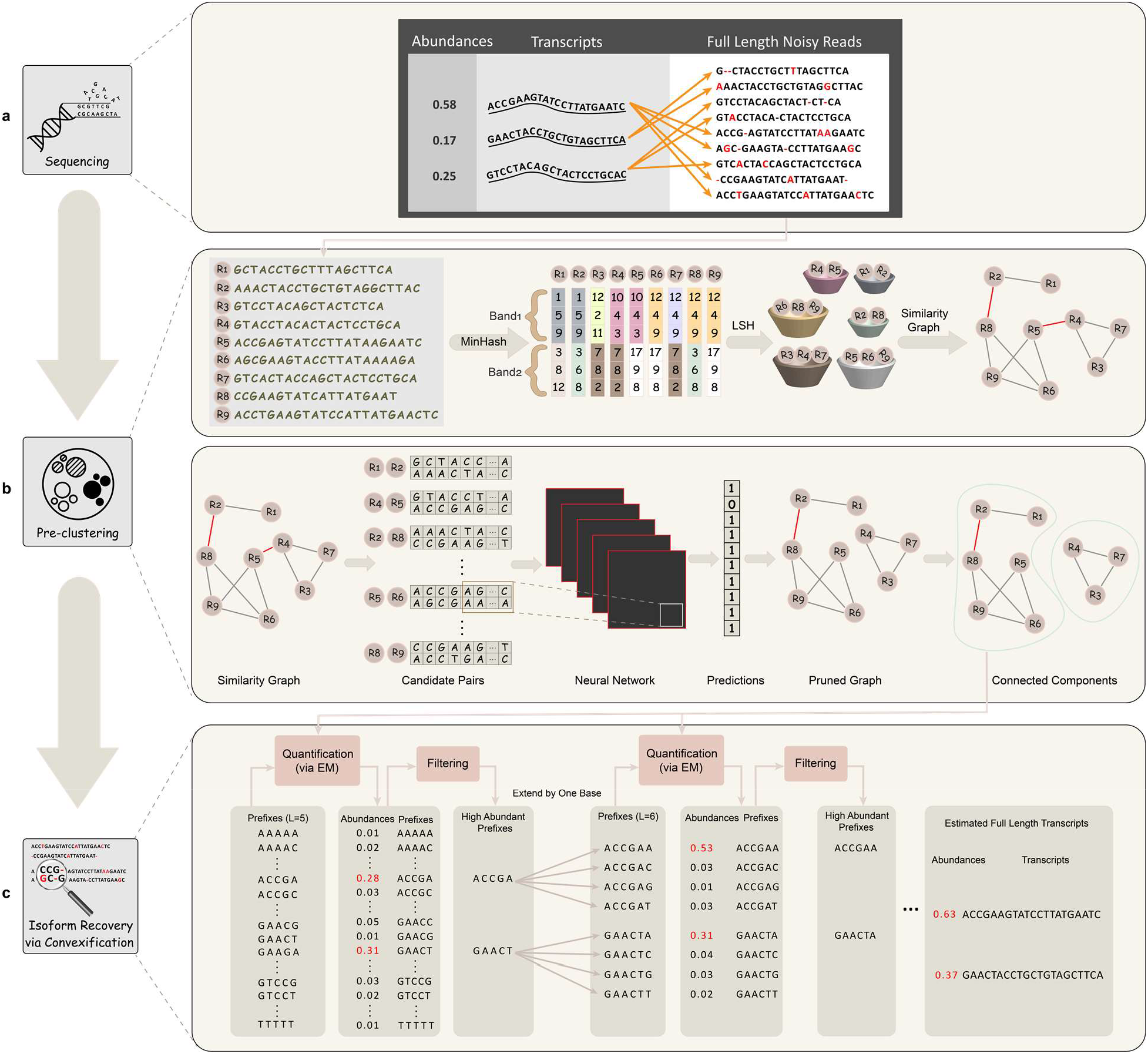
I-CONVEX algorithm workflow: **(a)** *Sequencing full-length transcriptome using long reads*. The reads cover the full transcripts and the number of reads from each transcript is proportional to its abundance in the sequencing library. Different error types such as insertion, deletion, and substitution may occur in the reads. The reads are the input to I-CONVEX. **(b)** *Pre-clustering stage:* First, we use MinHash to obtain the signature matrix. Then, the signature matrix is divided into several bands (e.g., two bands with size 3). Two reads are in the same bucket if they are equal in all rows of a band (e.g. R4, R5). Next, a similarity graph is formed by connecting reads that are in the same bucket. The edges of this graph are then validated with a neural network to reduce the number of false positive edges (red edges). Each connected component of the similarity graph leads to one pre-cluster. **(c)** An example run of *Clustering via Convexification*. First, the list of all possible short prefixes is considered (e.g. 4^5^ = 1024 prefixes of length **L = 5** ranging from ‘AAAAA’ to ‘TTTTT’). The abundances of these prefixes are then estimated by aligning them to the reads and solving a maximum likelihood estimation problem through the (sparse) expectation maximization (EM) algorithm^13,14^ (See Online methods). I-Convex only keeps the prefixes with the abundance higher than a specified threshold. Then each length **L** prefix ‘XXXXX’ is replaced by four extended prefixes ‘XXXXXA’, ‘XXXXXC’, ‘XXXXXG’, and ‘XXXXXT’. Using the previous alignment of the prefixes to the reads, the abundance of these length L+1 prefixes are estimated and the list is filtered and extended to obtain a list of prefixes of length L+2. This procedure continues until the complete recovery of all transcripts.

In contrast to ICE and IsoCon, our method I-CONVEX does not require read-to-read alignment. I-CONVEX consists of two subprograms: scalable pre-clustering of reads (**Figure 1b**), and alignment-free isoform recovery via convexification (**Figure 1c**). We first describe the alignment-free isoform recovery step (**Figure 1c**), which is the core module of I-CONVEX and is based on the following observation: When the list of transcripts is known, estimating the abundances is a convex problem and can be done efficiently using convex optimization approaches such as the EM algorithm^12,13^. However, the list of transcripts is not known in ***de novo*** transcriptome recovery *a priori*, which makes the problem non-convex. A convex reformulation of the problem could be obtained by assuming that all sequences are possibly transcripts (with many of the sequences having zero abundances). However, this reformulation would grow exponentially with sequence lengths. To overcome this exponential increase, we first reduce the size of the problem by partitioning the reads into a small number of equivalence classes that share the same (short) prefixes, then we estimate their aggregate abundances (**Figure 1c**). Many of the equivalence classes would have near-zero abundances that are then “pruned”. Keeping only the classes with sufficiently large abundance estimates, we further partition, or “branch” them by extending the prefixes one base at a time until a maximum length threshold is reached. At each step of the algorithm, the abundance of each equivalent class can be estimated using the EM algorithm with added sparsity regularization (**Supplementary Note 2**). The computational complexity of the algorithm grows linearly with the number of reads. This alignment-free isoform recovery step can fully utilize multiple computational cores by processing the reads in parallel (**Online Methods**). The parallelization is achieved without losing any statistical accuracy as the parallel version, and the single-core version returns exactly the same output.

To scale I-CONVEX to millions of reads, the first step of I-CONVEX performs a fast pre-clustering algorithm on the input reads by constructing a “conservative” similarity graph (**Figure 1b**). The nodes in this graph correspond to the reads, and an edge shows a similarity level higher than a certain threshold. A low threshold is chosen to capture any potential similarity among the reads. Thus, each connected component (pre-cluster) in this graph contains all the reads coming from a group of similar transcripts. To obtain the similarity graph, we use a locality sensitive hashing (LSH) method based on the Jaccard similarity^14–16^ between k-mer signatures. This idea has been used before in Mash^15^ and MHAP^16^. However, to make the computational complexity of the algorithm linear in the number of reads, we adopt the idea of banding technique^17^ (**Supplementary Note 3**). In this pipeline, the resulting similarity graph may contain a large number of false positive edges since k-mer sharing amongst non-homologous transcripts is frequent. To reduce the number of false positives, we trained a convolutional neural network to validate and correct the similarity of read pairs (**Supplementary Note 4**). Then, the obtained pre-clusters are processed in parallel by the clustering via convexification step (**Figure 1c**).

We compare the performance and efficiency of I-CONVEX against IsoCon and ICE on simulated and real datasets in **Figure 2** (see **Supplementary Note 5** for the details of the datasets). As can be seen in **Figure 2a**, I-CONVEX can efficiently scale to large size datasets. **Figure 2b** and **Figure 2d** compare the recall, precision, and F-score for I-CONVEX, ICE, and IsoCon on SIRV and simulated datasets. In contrast to ICE and IsoCon, the number of false positives in I-CONVEX output decreases as the number of reads increases. Thus, the F-score of I-CONVEX is enhanced by increasing the number of reads, while the other two methods suffer from low F-scores when we increase the sequencing coverage. We further evaluate I-CONVEX on the Sequel II dataset (**Supplementary Note 5**) containing approximately 7 million reads. Since the actual transcriptome is not available, we apply SQANTI2^18^ to the predicted transcriptome. SQANTI2 outputs the number and percentage of full-splice matches (perfect matches to a reference transcript), incomplete-splice matches (possible degraded matches to a reference transcript), and novel transcriptome (which are not high-quality transcripts with high probability) in the predicted transcripts. As depicted in **Figure 2c**, I-CONVEX generates fewer transcripts (high precision) compared to the IsoSeq3 (a successor of ICE in the PacBio SMRTAnalysis software suite), while the majority of them are either full-splice matches or incomplete-splice matches.

**Figure 2:**
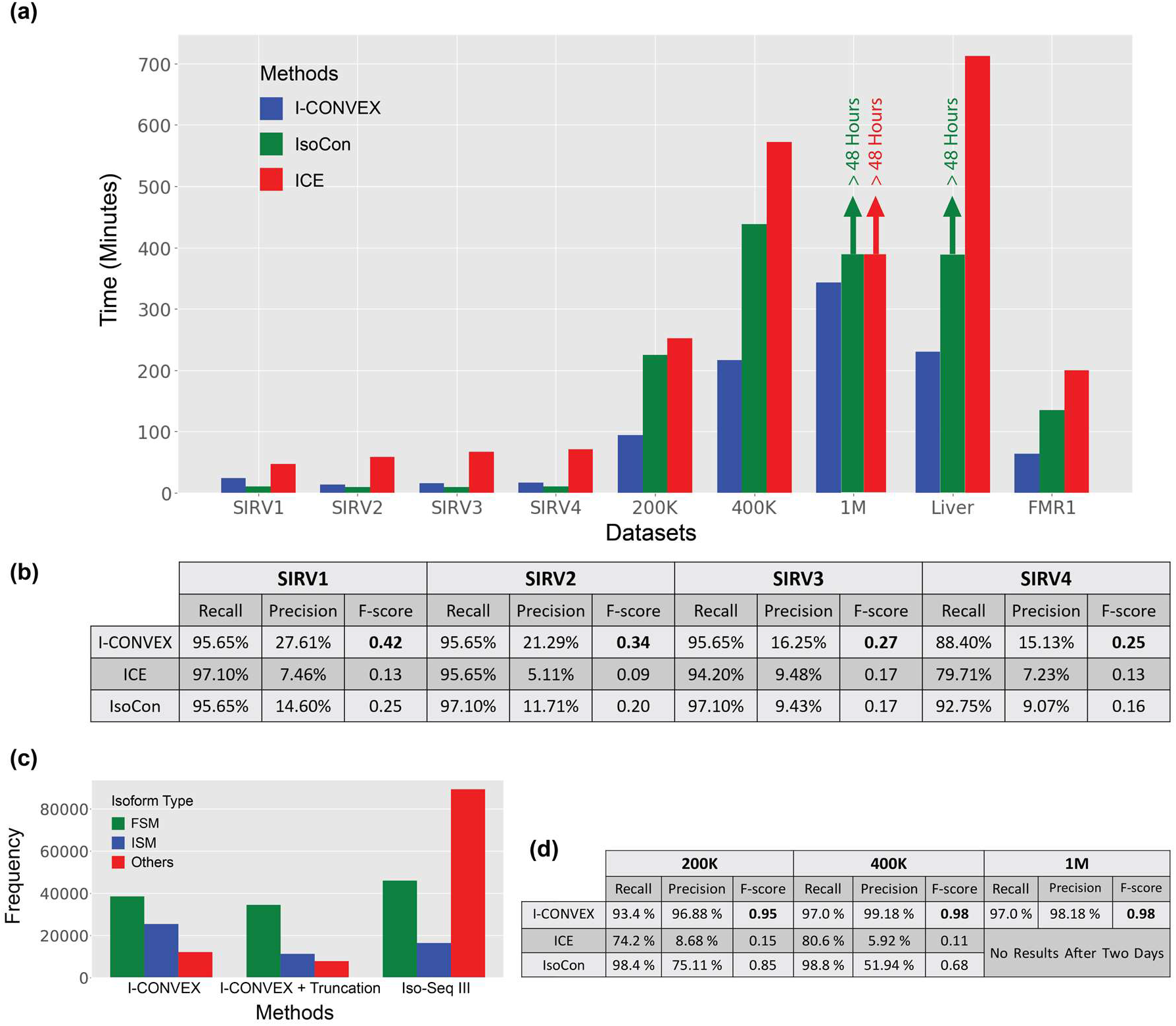
Performance Comparison of I-CONVEX, ICE, and IsoCon. **(a)** Running time of I-CONVEX, ICE, and IsoCon for real and simulated read datasets. The insertion/deletion/substitution error is generated according to the identical independent error model. All the methods have the same amount of memory (180 GB) and computational resources (16 cores per cluster). ICE and IsoCon could not complete the denoising task within 48 hours for the 1M dataset. In addition, IsoCon cannot complete the denoising task for the Liver dataset within 48 hours. **(b,d)** Comparison of the recall, precision, and F-score performed on several simulated datasets with known ground-truth. Recall measures the ratio of the actual transcripts detected with an accuracy of larger than 98%. Precision measures the number of recovered ground truth transcripts divided by the total number of estimated transcripts. While the recall of three methods is close to each other, I-CONVEX demonstrates a better performance in terms of precision. **(c)** The frequency of Full Splice Matches (FSM) and Incomplete Splice Matches (ISM) obtained by running Iso-Seq 3 and I-CONVEX on the Sequel II dataset. The “I-CONVEX + Truncation” means pre-clusters with size 1 are thrown away. We could not get the result of IsoCon and ICE on this dataset after waiting for more than 48 hours.

From a broader viewpoint, I-CONVEX solves a clustering problem over finite-alphabet sequences. The ability of I-CONVEX for fast and accurate clustering of the sequences can be beneficial in various other applications. For example, the read or k-mer denoising problems can be viewed as a clustering problem where reads/k-mers from identical sequences belong to the same cluster. As another example, the reconstruction of antibody repertoire, which is an important step in immunology and drug development, can be viewed as a clustering problem^19^ and the idea behind I-CONVEX could lead to linear time algorithms for this purpose.

The I-CONVEX package is available online at https://github.com/sinaBaharlouei/I-CONVEX. We have provided the basic instructions to run I-CONVEX in **Supplementary Note 6**.

## Methods

Methods and any associated references are available in the online version of the paper.

## Author Contributions

M.R and D.T proposed the clustering via convexification procedure of I-CONVEX (**Figure 1c**); M.R. implemented the clustering via convexification process (**Figure 1c**). S.B designed and implemented the pre-clustering process (**Figure 1b**) and tuned it with the convexification procedure. S.B and E.T tested the algorithm on different datasets and analyzed the results. All authors contributed in writing the manuscript.

## Methods

### Computing Abundances

The likelihood of observing the set of reads *R* = {*r*_1_, *r*_2_,…, *r_n_*} from a given set of transcripts *T* = {*t*_1_, *t*_2_,…, *t_n_*} with abundances *ρ* = {*ρ*_1_,…, *ρ_m_*} can be computed as^18^

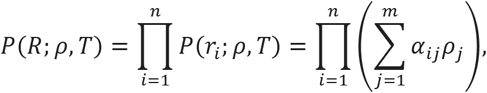

where *α_ij_* is the probability of observing the read *r_i_* from transcript *t_j_*. Therefore, the maximum likelihood estimation of *ρ* is given by

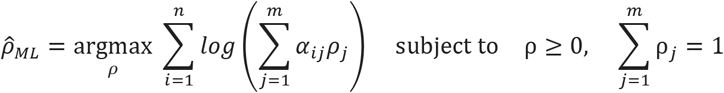

which can be solved through Expectation Maximization (EM) algorithm iteration^18^:

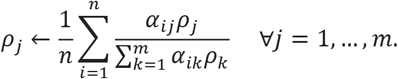

### Sparsification of the abundance vector estimation

The abundance of the sequences in the *Isoform Recovery via Convexification step* in I-CONVEX is a sparse vector. Hence, Isoform recovery via convexification step (**Figure 1c**) estimates the abundance vector through *l_q_*-norm regularization for imposing sparsity by solving

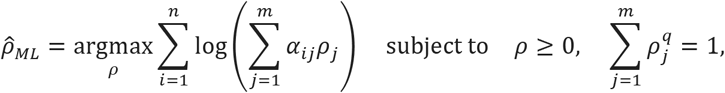

where *q* is some positive constant less than 1. In our experiments, we observe that setting the value of *q* close to one, e.g. 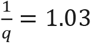, reduces the number of false positives while does not decrease the number of true positives. This modified optimization problem can be solved through the following iterative procedure (**Supplementary Note 2**):

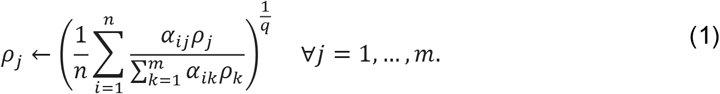

### Parallelization

Isoform recovery via convexification step (**Figure 1c**) partitions the reads evenly among the cores before running the algorithm. Each core keeps a copy of the estimated prefixes and abundances while it computes the parameters *α_ij_* for its own reads. Let us assume that the set of reads *R* = (*r*_1_,…, *r_n_*} is partitioned into subsets *R*_1_,…, *R_C_* with *c* being the number of computational cores. At each iteration of the algorithm, each core *l* computes local values

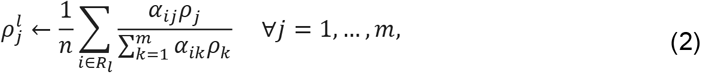

and then the consensus abundance value is obtained by

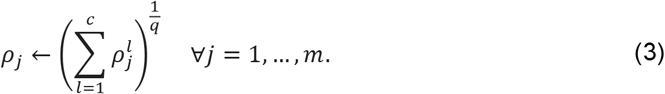

The above two steps (2, 3) are equivalent to (1) and return the exact same values for abundances.

### Pre-clustering

To reduce the time and memory complexity of the isoform recovery via convexification procedure, we propose a fast pre-clustering algorithm (**Figure 1b**) consisting of three main steps:

1. Fast mapping of reads to buckets based on MinHash and Locality Sensitive Hashing (LSH) algorithms. The more similar a pair of reads, the higher the probability of mapping them to the same bucket is.
2. Validating similar candidate pairs with a trained convolutional neural to eliminate false positive pairs obtained from the previous step.
3. Pre-clustering the similarity graph whose vertices are reads and edges show the similarity between the reads.

Each of these steps is explained in details below:

### Fast mapping of reads to buckets

To measure the proximity of read pairs, a widely used idea is to compute the Jaccard similarity of their k-mer set^16,17^. The k-mer set of a given read is the set of all of its consecutive subsequences with length *k*. As an example “GCTACCT” consists of {“GCTA”, “CTAC”, “TACC”, “ACCT”} 4-mers. For a given dataset containing *N* reads with the average length of *L*, it takes *O*(*NL*) operations to obtain the k-mer representation of all the reads. For convenience, each k-mer is hashed to a 32-bit integer number. Having the k-mer set of all the reads in the dataset, we form a representation matrix *M* with its columns representing different reads and different rows representing different k-mers (that appear in at least one read). Each entry *M_ij_* equals to 1 if and only if the *i*-th k-mer appears in the j-th read, and 0 otherwise. Since computing the Jaccard similarity of read pairs is computationally expensive, we compress the reads using MinHash signatures, which are unbiased estimators of Jaccard similarity (**Supplementary Theorem 2 in Supplementary Note 3**). Thus, instead of exact computation of Jaccard similarity of all read pairs, we can estimate them by finding the Hamming similarity of their MinHash signatures. *h* ≪ *L* MinHash functions are applied to the representation matrix *M* to obtain a MinHash signature with length *h* for each read. Choosing a larger value for *h*, corresponds to a smaller variance of the Jaccard estimator (**Supplementary Theorem 3 in Supplementary Note 3**). Hence a signature matrix S with *N* columns and h rows can be formed such that ¾ represents the i-th element of the MinHash code corresponding to the j-th read in the dataset. To compute *S_ij_*, let 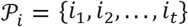 be a permutation of {1, 2,…, *t*} corresponding to the i-th MinHash function, where *t* is the number of rows in *M*. Let *i_min_* be the smallest integer such that *M*[*i_min_*] [*j*] = 1. Then, the MinHash value of the j-th read with respect to the permutation *P_i_* equals *i_min_*.

Computing the similarity of two MinHash signatures rather than the original k-mer sets is significantly more efficient. However, even after hashing long reads to MinHash signatures, calculating the similarity of all 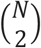 pairs of MinHash signatures is still a computationally expensive task. To avoid pairwise comparison of all reads, we adopt the locality sensitive hashing (LSH) algorithm. The corresponding MinHash signature of each read is divided into *b* bands with size *d* (*h* = *bd*). Accordingly, the first *d* rows of the signature matrix form the first band. The second *d* rows of the signature matrix correspond to the second band, and so on. If two columns (which correspond to two different reads) are equal in all rows of at least one of these bands, they will be mapped to the same bucket, and we call them a **candidate similar pair** (**Figure 1b**). Assume s is the true Jaccard similarity of *S*_1_ and *S*_2_. Since MinHash is an unbiased estimator of the Jaccard similarity, the probability that *S*_1_ and *S*_2_ are equal in each row is *s*. Thus, the probability of being equal in all *d* rows of a band is *s^d^*. Hence *S*_1_ and *S*_2_ will be mapped to the same bucket with the probability *p* = 1 – (1 – *s^d^*)^*b*^. *d* and *b* can be seen as two hyper-parameters that control the false positive and false negative rates. Increasing *d* leads to the decrease in the value of *p*. Thus, the number of pairs mapped to the same bucket is decreased; and true positive and false positive rates are reduced simultaneously (**Supplementary Figure 2**). The same logic implies that by increasing the number of bands (*b*), *p* will be increased. Therefore, both true positive and false positive rates will be increased. To avoid low true positive rate, we choose small values for *d* and *b*, and then we eliminate the false similar candidate pairs using a trained convolutional neural network.

### Validating Candidate Similar Pairs via Convolutional Neural Networks

To validate the candidate pairs obtained by applying LSH on the dataset of reads, we designed a Convolutional Neural Network (CNN), which takes a pair of sequences and generates the output one if the sequences are similar and zero otherwise (**Figure 1b**). **Supplementary Table 1** depicts the architecture of the designed convolutional neural network in detail. The training data consists of 100000 pairs of the reads, where half of them are similar, and the rest are dissimilar (The details of the training dataset is available in **Supplementary Note 4**). We optimize the following objective function applying an Adam optimizer with the step-size *α* = 10^-4^ and the momentum *β*_1_ = 0.9:

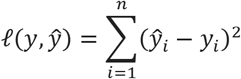

### Pre-clustering the Similarity Graph

The similarity graph among the reads is an undirected graph in which a vertex represents a read in the dataset. We connect two vertices with an edge if and only if their corresponding reads are detected as a similar pair by the convolutional neural network introduced in the previous step. Ideally, if the LSH algorithm combined with the validation phase by the designed CNN can detect all the similar pairs without producing any false edges, each connected component of the similarity graph corresponds to one cluster. However, due to the existence of false positives in the graph, each connected component may contain more than one actual cluster. In practice, there is typically a large connected component containing 10% to 40% of the nodes. For this specific component, we run a fast greedy community detection algortihm^20^ to find the pre-clusters it contains.

Partitioning the reads into pre-clusters has several advantages over running the “isoform recovery via convexification” module on the entire dataset. First, this module can be executed on different pre-clusters independently and in parallel. Hence, the amount of memory per task, and the entire time needed for the transcriptome recovery will be decreased profoundly as a result of parallelization. Second, the time complexity of the convexification stage is dependent on the product of the number of transcripts and the number of reads. Since the number of transcripts in each pre-cluster is much smaller than the total number of transcripts, pre-clustering significantly improves the computational complexity of “isoform recovery via convexification” step.

