## Supplementary Notes for "I-CONVEX: Fast and Accurate *de Novo* Transcriptome Recovery from Long Reads"

**Supplementary Note 1: Identifiability of *de novo* Transcriptome Recovery from Long Reads**

A family of distributions $P = \{P_\theta | \theta \in \Theta\}$ parameterized by a vector $\theta$ is called identifiable if it satisfies the following condition:

$$P_{\theta_1} = P_{\theta_2} \Longrightarrow \theta_1 = \theta_2 \quad \forall \theta_1, \theta_2 \in \Theta,$$

This means that with enough observations from one of the distributions inside the family, one can decide which $\theta$ is the ground-truth value. For the *de novo* transcriptome recovery problem, $\theta$ is defined as the set of unknown transcripts/sequences $S = \{s_1, s_2, \ldots, s_m\}$ and their corresponding abundances $\rho = \{\rho_1, \rho_2, \ldots, \rho_m\}$. For a given read $r$, we define $P_\theta(r)$ as the probability of observing $r$, given $\theta = (S, \rho)$. We assume that a read generated from a sequence by adding insertion, deletion, or substitution errors through an i.i.d. process which does not depend on the location of the error. For simplicity, let $\delta_s$, $\delta_i$, and $\delta_d$ be the probability of observing a substitution, insertion, or deletion error at each location of the sequence respectively. In **Theorem 1**, we prove that the problem of *de novo* transcriptome recovery is identifiable.

**Supplementary Theorem 1 (Long-read Sequencing Identifiability):** Assume that the set of sequences $S = \{s_1, s_2, \ldots, s_m\}$ with the corresponding abundances $\rho = \{\rho_1, \rho_2, \ldots, \rho_m\}$ is unknown. Suppose that for any read $r$, $P_{S,\rho}(r)$ represents the probability of observing read $r$ given that it is generated from a sequence in $S$ with the abundance vector $\rho$. Then, given $P_{S,\rho}(r)$ for all reads $r$, both $S$ and $\rho$ can be exactly recovered when substitution error parameter $\delta_s \neq \frac{3}{4}$.

**Proof:** Each read $r$ with non-zero $P(r)$ can be obtained by adding insertions, deletions, and substitutions to one or several sequences in $S$. Without loss of generality, we can model the transcriptome recovery problem as denoising the reads obtained by applying three noise channels to the set of sequences in $S$ sequentially (**Supplementary Figure 1**). In this model, $P_1(r)$ is the distribution of reads after the substitution channel; and $P_2(r)$ is the distribution of reads after applying the deletion channel (and before the insertion channel). Lastly, $P(r)$ is the final probability distribution observed after applying the insertion noise. If we prove that the transcriptome recovery is identifiable under each one of these three channels (insertion, deletion, and substitution) individually, then the entire problem is identifiable. This is because of the fact that using the given probability of observing reads $P(r)$, we can recover $P_2(r)$ by denoising the effect of the insertion channel. Then, with the same argument, we can recover $P_1(r)$ by denoising the effect of the deletion channel. Finally, we can find the vector $\rho$ by denoising the effect of the substitution channel on the probability vector $P_1$. Thus, it suffices to show that $P_2(r)$, $P_1(r)$, and $\rho$ can be recovered sequentially given $P(r)$. In what follows, we show each one of these three channels is identifiable.

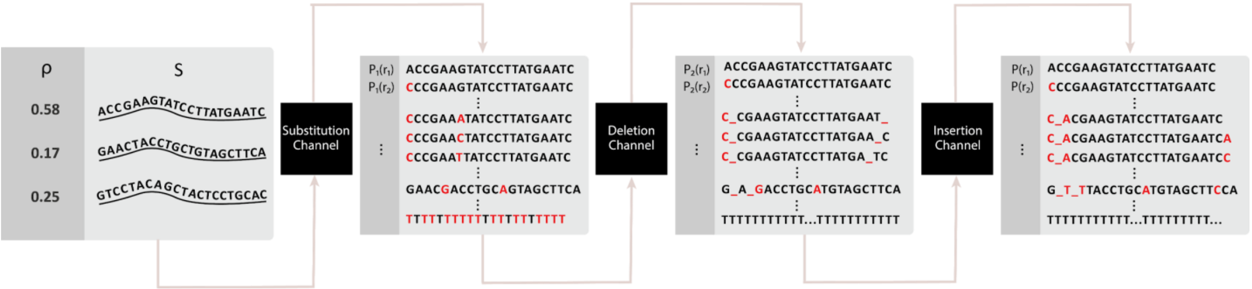


**Supplementary Figure 1:** Each channel affects the probability vector of reads from the previous step. To prove the identifiability of the transcriptome recovery problem, it suffices to denoise the effect of the channels sequentially from last to first.

**Identifiability under the substitution channel:** For the substitution channel, we can observe that a read $r$ with length $L$ can only be generated from a sequence of size $L$, since the substitution does not change the sequence length. Thus, for estimating the abundances of the sequence with size $L$, we can only focus on the set of reads with size $L$. It means that without loss of generality we can assume that the set of reads and sequences are all of size $L$. Moreover, without loss of generality, assume that S consists of all $4^L$ possible transcripts with size $L$ with possibly zero abundances. Thus, the problem is reduced to recovering vector $\rho$, given the $P(r)$ and $S$. Let $\delta_s = \delta$ be the substitution error rate. Then the probability of observing the read $r_i$ can be written as follows:

$$P(r_i) = \sum_{j=1}^{n} \rho_j P(r_i|s_j) = \sum_{j=1}^{n} \rho_j \left(\frac{\delta}{3}\right)^{d(r_i,s_j)} (1-\delta)^{L-d(r_i,s_j)}$$

Here $d(r_i, s_j)$ is the number of mismatches between $r_i$ and $s_j$. This relationship can be written in a matrix form. For example, when $L = 1$, let the abundances of $'A', 'C', 'G'$ and $'T'$ be $\rho_1, \rho_2, \rho_3$ and $\rho_4$ respectively. Then, the probability of observing each $\{'A', 'C', 'G', 'T'\}$ can be obtained by the following equation:

$$\begin{bmatrix} P(r_1) \\ P(r_2) \\ P(r_3) \\ P(r_4) \end{bmatrix} = A \begin{bmatrix} \rho_1 \\ \rho_2 \\ \rho_3 \\ \rho_4 \end{bmatrix}$$

where

$$A = \begin{bmatrix} 1-\delta & \delta/3 & \delta/3 & \delta/3 \\ \delta/3 & 1-\delta & \delta/3 & \delta/3 \\ \delta/3 & \delta/3 & 1-\delta & \delta/3 \\ \delta/3 & \delta/3 & \delta/3 & 1-\delta \end{bmatrix}$$

Thus, the identifiability problem in this case reduces to showing the invertibility of this matrix. Notice that the determinant of this matrix can be computed as

$$\begin{vmatrix} 1-\delta & \delta/3 & \delta/3 & \delta/3 \\ \delta/3 & 1-\delta & \delta/3 & \delta/3 \\ \delta/3 & \delta/3 & 1-\delta & \delta/3 \\ \delta/3 & \delta/3 & \delta/3 & 1-\delta \end{vmatrix}$$
$$= (1-\delta) \begin{vmatrix} 1-\delta & \delta/3 & \delta/3 \\ \delta/3 & 1-\delta & \delta/3 \\ \delta/3 & \delta/3 & 1-\delta \end{vmatrix} - \frac{\delta}{3} \begin{vmatrix} \delta/3 & \delta/3 & \delta/3 \\ \delta/3 & 1-\delta & \delta/3 \\ \delta/3 & \delta/3 & 1-\delta \end{vmatrix} + \frac{\delta}{3} \begin{vmatrix} \delta/3 & 1-\delta & \delta/3 \\ \delta/3 & \delta/3 & \delta/3 \\ \delta/3 & \delta/3 & 1-\delta \end{vmatrix}$$
$$- \frac{\delta}{3} \begin{vmatrix} \delta/3 & 1-\delta & \delta/3 \\ \delta/3 & \delta/3 & 1-\delta \\ \delta/3 & \delta/3 & \delta/3 \end{vmatrix} = (1-\delta) \begin{vmatrix} 1-\delta & \delta/3 & \delta/3 \\ \delta/3 & 1-\delta & \delta/3 \\ \delta/3 & \delta/3 & 1-\delta \end{vmatrix} - \delta \begin{vmatrix} \delta/3 & \delta/3 & \delta/3 \\ \delta/3 & 1-\delta & \delta/3 \\ \delta/3 & \delta/3 & 1-\delta \end{vmatrix}$$
$$= (1-\delta)\left(\frac{8}{27}\delta^3 + \frac{3}{2}\delta^2 - 2\delta + 1\right) - \delta\left(\frac{16}{27}\delta^3 - \frac{8}{9}\delta^2 + \frac{\delta}{3}\right) = \frac{24}{27}\delta^4 + \frac{14}{27}\delta^3 + \frac{7}{3}\delta^2 - 3\delta + 1,$$

where the second equality is due to the effect of permuting rows on the determinant of the matrix. One can easily check that this polynomial is non-zero for all $\delta \in [0, 1]$ except $\delta = \frac{3}{4}$, which implies the invertibility of the matrix for all $\delta \in [0, 1]$ except $\delta = \frac{3}{4}$.

Now, let us consider the case of general length $L > 1$. Assume that we order the set of all possible sequences with size $L$ (all $m = 4^L$ possible sequences) according to the order of sequences in base $4$. In this case, the probability transition relation can be written as

$$\begin{bmatrix} P(r_1) \\ P(r_2) \\ \dots \\ P(r_m) \end{bmatrix} = \underbrace{A \otimes \dots \otimes A}_{L \text{ times}} \begin{bmatrix} \rho_1 \\ \rho_2 \\ \dots \\ \rho_m \end{bmatrix}$$

where the Kronecker product of two matrices $A_{(m \times n)}$ and $B_{(p \times q)}$ is a matrix $C_{((m\times p) \times (n\times q))}$ , defined as:

$$C = A \otimes B = \begin{bmatrix} a_{11}B & a_{12}B & \dots & a_{1n}B \\ a_{21}B & a_{22}B & \dots & a_{2n}B \\ \dots & \dots & \dots & \dots \\ a_{m1}B & a_{m2}B & \dots & a_{mn}B \end{bmatrix}$$

The Kronecker product of $L$ matrices is invertible if and only if each one of them is invertible[21]. We have already shown that $A$ is an invertible matrix when $\delta_s \neq \frac{3}{4}$. Thus, $A \otimes A \otimes \dots \otimes A$ is invertible as well, which shows the problem is identifiable when the substitution channel is applied.

**Insertion Channel:** Let $R$ be the support of $P$ ($R = \{r: P(r) > 0\}$). We define the operator $\leq$ on a pair of reads as follows: $r_1 \leq r_2$ if $r_2$ can be obtained by inserting zero or more bases to $r_1$ in different locations. It is easy to observe that $(R, \leq)$ forms a partially ordered set (poset). Since it satisfies the reflexivity, anti-symmetry, and transitivity. We have the following observations:

- If $r' \leq r$ and $r \neq r'$, then $|r'| < |r|$.
- $R$ has at least one minimal element.
- Any minimal element of $R$ belongs to $S$.

To see the correctness of the first observation, assume that $r' \leq r$, which means r is obtained by adding zero or more characters to r. However, since $r' \neq r$, the number of added characters cannot be zero. Thus at least one character is added to $r'$ to obtain r. This means that $|r'| \leq |r|$.

To prove the second observation, assume that $r_{min}$ is an element of $R$ with the minimum length (it is not necessarily unique). Indeed, such an element exists, because $|r| \geq 1$ for all $r \in R$. We claim that $r_{min}$ is a minimal element. We prove by contradiction. Assume that $r_{min}$ is not a minimal element. Thus, there exists an element $r' \neq r_{min}$ in $R$ such that $r' \leq r_{min}$. According to the first observation $|r| < |r_{min}|$, which contradicts the fact that $r_{min}$ is an element with the minimum length in $R$.

Finally, we prove that a minimal element of $R$ belongs to $S$. We prove by contradiction. Assume that $r_{min}$ is a minimal element that does not belong to $S$. Thus, there exists an element $s \in S$, such that $s \leq r_{min}$. Clearly $r = s$ belongs to $R$, since $P(r) \geq P(r|s) = \rho_s(1 - \delta_i)^{|s|} > 0$. Therefore, $s$ is an element in $R$ which is not equal to $r_{min}$ and $s \leq r_{min}$ which contradicts with the fact that $r_{min}$ is a minimal element in $R$.

Now, we prove the identifiability of the transcriptome recovery problem under the insertion channel by induction on the size of $S$.

For $|S| = 1$, according to the third observation, the minimal member of $R$ belongs to $S$, and its abundance is clearly $1$.

Based on the induction hypothesis, assume that the problem is identifiable when $|S| = m$. We want to prove it is identifiable, when $|S| = m + 1$. Based on the third observation, any minimal element $r_{min}$ of $R$ belongs to $S$. Set $S' = S - \{r_{min}\}$ and update the probability vector of the reads accordingly: $P'(r) = P(r) - P(r|r_{min})\, \rho_{r_{min}}$. For updating the probability vector of reads we need to estimate $\rho_{r_{min}}$ exactly. Since $r_{min}$ is the minimal element, it can be only obtained from a sequence with the exact same structure. Hence, $P(r_{min}) = \rho_{r_{min}}(1-\delta)^{|r_{min}|}$, which means $\rho_{r_{min}} = \frac{P(r_{min})}{(1-\delta)^{|r_{min}|}}$. According to the assumption of induction, since $|S'| = m$, we can recover $S'$ using $P'$. Thus, $S = S' \cup \{r_{min}\}$, which means we can recover $S$ exactly.

**Deletion Channel:** The argument is similar to the insertion channel. The only difference is that the operator $\leq$ is defined for a pair of reads $r_1 \leq r_2$ if $r_2$ can be obtained by deleting some elements of $r_1$. Moreover, the minimal element of $R$ is the longest read in this case.

**Supplementary Note 2: Sparse EM Algorithm Inside the Isoform Recovery via Convexification**

As discussed in Section **Methods**, the maximum likelihood estimator of the abundance vector given the set of transcripts and reads can be written as:

$$\hat{\rho}_{ML} = \underset{\rho}{\text{argmax}} \sum_{i=1}^{n} log\left(\sum_{j=1}^{m} \alpha_{ij}\rho_j\right) \quad \text{subject to} \quad \rho \geq 0, \quad \sum_{j=1}^{m} \rho_j = 1$$

Or equivalently:

$$\hat{\rho}_{ML} = \underset{\rho}{\text{argmin}} \sum_{i=1}^{n} -log\left(\sum_{j=1}^{m} \alpha_{ij}\rho_j\right) \quad \text{subject to} \quad \rho \geq 0, \quad \sum_{j=1}^{m} \rho_j = 1$$

Applying the Expectation Maximization (EM) algorithm, at each iteration, we minimize a tight local upper bound of the above objective function:

$$\rho^{t+1} = \underset{\rho}{\text{argmin}} - \sum_{i=1}^{n}\sum_{j=1}^{m} \frac{\alpha_{ij}\rho_j^t}{\sum_{j'=1}^{m}\alpha_{ij'}\rho_{j'}^t} \log\left(\frac{\rho_j}{\rho_j^t}\right) - \sum_{i=1}^{n} \log\left(\sum_{j=1}^{m} \alpha_{ij}\rho_j\right) \quad \text{subject to} \quad \rho \geq 0, \quad \sum_{j=1}^{m} \rho_j = 1$$

The Lagrangian function of the above problem can be written as:

$$L(\rho, \lambda) = - \sum_{i=1}^{n}\sum_{j=1}^{m} \frac{\alpha_{ij}\rho_j^t}{\sum_{j'=1}^{m}\alpha_{ij'}\rho_{j'}^t} \log\left(\frac{\rho_j}{\rho_j^t}\right) - \sum_{i=1}^{n} \log\left(\sum_{j=1}^{m} \alpha_{ij}\rho_j\right) + \lambda\left(\sum_{j=1}^{m} \rho_j - 1\right)$$

And the dual problem takes the form of:

$$\max_{\lambda} \min_{\rho} L(\rho, \lambda)$$

Since the Lagrangian function is convex in $\rho$, and the constraints are linear, the strong duality holds[22]. Thus, by setting the gradient of $L$ with respect to $\rho$ to 0, we have:

$$- \sum_{i=1}^{n} \frac{\alpha_{ik}\rho_k^t}{\sum_{j'=1}^{m}\alpha_{ij'}\rho_{j'}^t} \frac{1}{\rho_k} + \lambda^* = 0, \quad \forall k = 1, \dots, m.$$

Which means:

$$\rho_k^{t+1} = \frac{1}{\lambda^*} \sum_{i=1}^{n} \frac{\alpha_{ik}\rho_k^t}{\sum_{j'=1}^{m}\alpha_{ij'}\rho_{j'}^t} \quad \forall k = 1, \dots, m.$$

Since $\sum_{j=1}^{m} \rho_j^{t+1} = 1$, we have:

$$1 = \sum_{k=1}^{m} \rho_k^{t+1} = \sum_{k=1}^{m} \frac{1}{\lambda^*} \sum_{i=1}^{n} \frac{\alpha_{ik}\rho_k^t}{\sum_{j'=1}^{m}\alpha_{ij'}\rho_{j'}^t} = \frac{1}{\lambda^*} \sum_{i=1}^{n} \frac{\sum_{k=1}^{m}\alpha_{ik}\rho_k^t}{\sum_{j'=1}^{m}\alpha_{ij'}\rho_{j'}^t} = \frac{n}{\lambda^*}$$

Therefore $\lambda^* = n$, and

$$\rho_k^{t+1} = \frac{1}{n} \sum_{i=1}^{n} \frac{\alpha_{ik}\rho_k^t}{\sum_{j'=1}^{m}\alpha_{ij'}\rho_{j'}^t} \quad \forall k = 1, \dots, m.$$

This update rule is similar to the one used by Express[12] to obtain the abundance vector.

**Sparsification.** One naïve approach to impose sparsity to the estimated abundances in the above problem is to use the $\ell_1$ regularizer[23]. However, the abundances are non-negative and their summation is equal to 1. Therefore, the $\ell_1$ regularizer does not change the objective function at all. Thus, instead of using $\ell_1$ regularizer, we apply the $\ell q$ regularizer to the objective function with $0 < q < 1$. Using this regularizer, the above problem can be written as

$$\rho^{t+1} = \underset{\rho}{\operatorname{argmin}} - \sum_{i=1}^{n} \sum_{j=1}^{m} \frac{\alpha_{ij}\rho_j^t}{\sum_{j'=1}^{m} \alpha_{ij'}\rho_{j'}^t} \log\left(\frac{\rho_j}{\rho_j^t}\right) - \sum_{i=1}^{n} \log\left(\sum_{j=1}^{m} \alpha_{ij}\rho_j\right) \text{ subject to } \quad \rho \geq 0, \quad \|\rho\|_q \leq 1$$

Thus, the Lagrangian function for the sparse EM takes the following form:

$$L(\rho, \lambda) = - \sum_{i=1}^{n} \sum_{j=1}^{m} \frac{\alpha_{ij}\rho_j^t}{\sum_{j'=1}^{m} \alpha_{ij'}\rho_{j'}^t} \log\left(\frac{\rho_j}{\rho_j^t}\right) - \sum_{i=1}^{n} \log\left(\sum_{j=1}^{m} \alpha_{ij}\rho_j\right) + \lambda\left(\left(\sum_{j=1}^{m} \rho_j^q\right)^{\frac{1}{q}} - 1\right)$$

Again, by writing the optimality condition for the dual problem, we have

$$\nabla L_{\rho}(\rho, \lambda) = 0 \ \Rightarrow$$
$$- \sum_{i=1}^{n} \frac{\alpha_{ik}\rho_k^t}{\sum_{j'=1}^{m} \alpha_{ij'}\rho_{j'}^t} \frac{1}{\rho_k^{t+1}} + \lambda^* q (\rho_k^{t+1})^{q-1} ((\rho_1^{t+1})^q + \cdots + (\rho_m^{t+1})^q)^{\frac{1}{q}-1} = 0, \quad \forall k = 1, \ldots, m.$$

Moreover, according to the complementary slackness, we have:

$$\lambda^* \left[\left(\sum_{j=1}^{m} \left(\rho_j^{t+1}\right)^q\right)^{\frac{1}{q}} - 1\right] = 0$$

It is easy to observe that $\lambda^* \neq 0$, otherwise the last two equalities have no solution. Thus, similar to the previous case, we can conclude that $\lambda^* = n$, and we have the following closed-form update rule:

$$\rho_k^{t+1} = \left(\frac{1}{n} \sum_{i=1}^{n} \frac{\alpha_{ik}\rho_k^t}{\sum_{j'=1}^{m} \alpha_{ij'}\rho_{j'}^t}\right)^{\frac{1}{q}}, \quad \forall k = 1, \ldots, m.$$

We refer to $\frac{1}{q}$ as the sparsity power. This hyper-parameter is set to $1.03$ in the implementation of *Isoform Recovery via Convexification* module. As we increase the sparsity power, the abundance vector is sparser, and thus the number of recovered isoforms is smaller at the end. Sparsification of EM can help to remove false positive isoforms further. However, we should avoid setting it to very large numbers as it may lead to the removal of true positives.

**Supplementary Note 3: Further Analysis of MinHash and Locality Sensitive Hashing Algorithms**

**Definition 1:** Suppose that $S_1$ and $S_2$ are k-mer sets of two given sequences. The Jaccard similarity of $S_1$ and $S_2$ is defined as

$$J(S_1, S_2) = \frac{|S_1 \cap S_2|}{|S_1 \cup S_2|}$$

**Definition 2:** Given a set of sequences $S_1, \dots, S_m$ and the set of k-mers $S'_1, \dots, S'_{N'}$, we say a matrix $M \in \{0,1\}^{N' \times m}$ is a representation matrix if it satisfies the following property: The element in row $i$ and column $j$ equals $1$ if and only if the i-th k-mer appears in j-th sequence, and $0$ otherwise.

**Definition 3:** Assume the representation matrix $M$ has $N'$ rows and $m$ columns, and let $\mathcal{P} = \{i_1, i_2, \dots, i_{N'}\}$ be a permutation of set $\{1, 2, \dots, N'\}$, and column $c$ in $M$ corresponds to a sequence $S$. Then the MinHash signature of $S$ with respect to $\mathcal{P}$ is defined as $\mathrm{MinHash}_{\mathcal{P}}(c) = i_j$ where $j$ is the smallest number in $\{1, 2, \dots, N'\}$ such that $M[i_j][c] = 1$.

**Example:** Assume that $S_1 = \mathrm{ACCAGTC}$ and $S_2 = \mathrm{ACCGTCA}$. The set of 3-mers that appears at least once in $S_1$ or $S_2$ is $\{\mathrm{ACC}, \mathrm{CCA}, \mathrm{CAG}, \mathrm{AGT}, \mathrm{GTC}, \mathrm{CCG}, \mathrm{CGT}, \mathrm{TCA}\}$. Let $\mathcal{P} = \{7, 5, 2, 3, 1, 4, 6\}$ be a random permutation of 3-mer indices. Since $5$ is the first 3-mer index in $\mathcal{P}$ such that its corresponding 3-mer appears in $S_1$, thus, the MinHash signature of $S_1$ with respect to $\mathcal{P}$ is $5$. With the same logic, we can observe that the MinHash signature of $S_2$ with respect to $\mathcal{P}$ is $7$.

**Supplementary Theorem 2:** Let $J(S_1, S_2)$ be the Jaccard similarity between two given sequences $S_1$ and $S_2$, and $\mathcal{P}_1, \dots, \mathcal{P}_h$ are $h$ distinct randomly generated permutations of set $\{1,2, \dots, N'\}$. For each sequence, a vector of MinHash signatures with length $h$ is computed with respect to permutations $\mathcal{P}_1, \dots, \mathcal{P}_h$. Then, $\frac{\#\text{Matches between signature vectors of } S_1 \text{ and } S_2}{h}$ is an unbiased estimator of the Jaccard similarity of $S_1$ and $S_2$.

**Proof:** Let $c_1$ and $c_2$ be the indices of columns in $M$ corresponding to $S_1$ and $S_2$. Let $R_1$ denote the set of rows in which $c_1$ and $c_2$ both have the value $1$, and $R_2$ be the set of rows in which exactly one of $c_1$ and $c_2$ is $1$. Fix a permutation $\mathcal{P} = \{i_1, i_2, \dots, i_{N'}\}$, and let $j$ be the smallest index such that at least one of $M[i_j][c_1]$ or $M[i_j][c_2]$ is $1$. Thus $i_j \in R_1 \cup R_2$. Moreover, $\mathrm{MinHash}_{\mathcal{P}}(\mathrm{c}_1) = \mathrm{MinHash}_{\mathcal{P}}(\mathrm{c}_2) = i_j$ if and only if $i_j \in R_1$. Hence $Pr[\mathrm{MinHash}_{\mathcal{P}}(\mathrm{c}_1) = \mathrm{MinHash}_{\mathcal{P}}(\mathrm{c}_2)] = \frac{|R_1|}{|R_1| + |R_2|}$ which is equal to the Jaccard similarity of $S_1$ and $S_2$. Thus, we have

$$\mathbb{E}\left[\frac{\#\text{Matches between signature vectors of } S_1 \text{ and } S_2}{h}\right] = \frac{\sum_{k=1}^{h} Pr\,[\mathrm{MinHash}_{\mathcal{P}_k}(\mathrm{c}_1) = \mathrm{MinHash}_{\mathcal{P}_k}(\mathrm{c}_2)]}{h}$$
$$= \frac{hJ(S_1, S_2)}{h} = J(S_1, S_2),$$

which means the estimator is unbiased.

**Supplementary Theorem 3:** The variance of the estimator proposed in **Theorem 2** is equal to $\frac{J(S_1,S_2) - J^2(S_1,S_2)}{h}$.

**Proof:** Since the probability of match between two MinHash signatures for each permutation $\mathcal{P}_k$ equals to $J(S_1, S_2)$, thus $\#\text{Matches between signature vectors of } S_1 \text{ and } S_2$ follows a binomial distribution with $h$ trials (number of permutations) and $p = J(S_1, S_2)$. Thus its variance equals $hp(1-p)$. Therefore,

$$\mathrm{Var}\left[\frac{\#\text{Matches between two signatures}}{h}\right] = \frac{\mathrm{Var}[\#\text{Matches between two signatures}]}{h^2} = \frac{hJ(S_1,S_2)\big(1-J(S_1,S_2)\big)}{h^2}$$
$$= \frac{J(S_1,S_2)\big(1-J(S_1,S_2)\big)}{h}$$

To obtain the MinHash signatures of reads in a given dataset, we need $O(NL)$ operations to form k-mer set of sequences, and $O(hNL)$ operations to obtain MinHash signatures of reads, where $N$ is the number of reads, $L$ is the average length of the sequences, and $h$ is the length of MinHash signatures.

### Choosing the hyper-parameters of the locality sensitive hashing Algorithm

Hashing the reads using MinHash reduces the computational complexity of comparison significantly, since comparing the MinHash signature of two given reads needs $O(h)$, while computing the edit distance of two reads requires $O(L^2)$ operations ($L$ can be varied from $400$ to $10k$, while $h$ can be selected as a constant number typically less than $100$). Still, comparing the MinHash signatures of all $O(N^2)$ pairs of sequences is inefficient for large-scale datasets consisting of millions of sequences. LSH algorithm copes with this issue by dividing the $h$ entries of the MinHash signature into $b$ bands of size $d$ $(h = bd)$. If two sequences are equal in all rows of at least one of the bands, they are considered as a candidate similar pair. For the default hyper-parameters of I-CONVEX $(d = 1,\ b = 10,\ k = 15)$ the probability of considering two sequences as similar for different values of their Jaccard similarity is depicted in **Supplementary Figure 2**. We have selected the default values of hyper-parameters such that the probability of having a false negative is small. However, this leads to a higher rate of false positives as well. As we mentioned in **Methods**, we validate the obtained candidate pairs by a designed convolutional neural network to remove the false positive edges in our similarity graph.

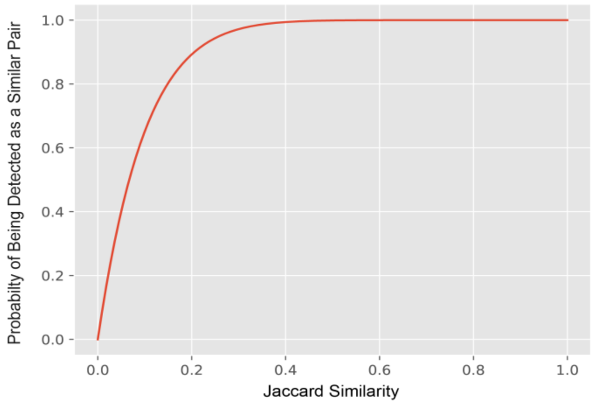


**Supplementary Figure 2:** The Probability of considering a pair of sequences as similar when d = 1 and b = 10 for different values of Jaccard similarity of the pair.

**Supplementary Note 4: Training the Neural Network for Validating Pairwise Similarities**

**Training Data:** To train the convolutional neural network validating the similarity of candidate similar read pairs, we have generated a balanced dataset of $50000$ similar and $50000$ dissimilar read pairs from a set of simulated transcripts. Two reads are labeled as similar if they are generated from a single transcript by adding insertion, deletion and substitutions. A naïve idea to simulate transcripts is to create completely random sequences consisting of {A, C, G, T}. However, in real datasets, it is highly probable that different transcripts have common sub-strings (exons). To address this issue, we first generate a pool of 400 exons $(e_1, e_2, \dots, e_{400})$. Each exon is a sequence of {A, C, G, T} with length $20$. Each simulated transcript is generated by concatenating $20$ exons $\{e_{i_1}, e_{i_2}, \dots, e_{i_{20}}\}$ picked from the pool in order $(i_1 < i_2 < \dots < i_{20})$. We repeat this procedure to simulate $100$ transcripts. To generate a similar pair, we choose a transcript and generate two reads by adding insertions, deletions, and shifts in the beginning and the end of the selected transcript. The insertion and deletion errors are both $2\%$. The length of the shift is a random variable, uniformly chosen from $0$ to $5$. To generate a dissimilar pair, we do the same, except that the reads are generated by perturbing two different transcripts. Compared to the scenario where the transcripts are simulated completely at random, this approach leads to a more powerful convolutional neural network with higher accuracy on real datasets.

**Architecture:** To train the convolutional neural network that validates the similarity of candidate pairs, for each pair in the generated training dataset (which can be similar or dissimilar), both reads are one-hot encoded first (Each letter corresponds to a $4 \times 1$ array). Since the length of each read in the generated dataset is 400, we obtain two arrays of size $4 \times 400$ after one-hot encoding of the reads. Then, we concatenate these two arrays row-wise to obtain an $8 \times 400$ array. These $8 \times 400$ arrays are the input data to the convolutional neural network with the layers depicted in **Supplementary Table 1**.

| Layer | Input | Filter Size | Filters | Stride | Activation |
|---|---|---|---|---|---|
| Convolution 1 | 8X400 | 8X8 | 32 | 1 | Relu |
| Max Pool 1 | 8X400X32 | 2X2 | - | 2 | - |
| Convolution 2 | 8X100X32 | 5X5 | 64 | 1 | Relu |
| Max Pool 2 | 8X100X64 | 2X2 | - | 2 | - |
| Fully connected 1 | 4X50X64 | | 50 | | Relu |
| Fully connected 2 | 50 | | 1 | | Linear |

**Supplementary Table 1:** Structure of the convolutional neural network designed to validate the similarity of candidate pairs.

The final value after passing each $8 \times 400$ input from the convolutional neural network is a scalar representing the predicted value. This scalar ranges between $-1$ and $1$, representing the similarity of a given pair. To compute the loss, let $y_i$ and $\hat{y}_i$ be the predicted value, and the true label corresponding to the i-th candidate pair. We use $\ell_2$ loss defined as follows:

$$\ell(y, \hat{y}) = \sum_{i=1}^{n} (\hat{y}_i - y_i)^2$$

To optimize the weight parameters in the network, an Adam optimizer is applied to the $\ell_2$ loss.

**Validation of Candidate Similar Pairs:** To validate a given candidate similar pair by the trained convolutional neural network, first each read is trimmed to a sequence with length $400$ (if the length of a read is less than $400$, we add enough base 'A' to the end of the sequence to reach the length of $400$). Each one of the two trimmed reads is one-hot encoded to get an array of size $4 \times 400$. By concatenating them, an array with the size $8 \times 400$ is obtained. Then, it will be passed through the convolutional neural network, and it gives a number within the range of $[-1, +1]$ . If the number is greater than $0$, we consider them as a similar pair.

**Supplementary Note 5: Test Datasets Description**

**Generating Simulated datasets with 200K, 400K, and 1M reads:** To create the simulated datasets in **Figure 2** with 200K, 400K, and 1M number of reads, we have selected the first 500 transcriptomes from the GENCODE dataset. Then, the transcript abundances are drawn from a log-normal distribution and normalized. **Supplementary Figure 3** shows the histogram of the transcript abundances for the 200K dataset. The reads are sampled based on the independent identical error model. Random shifts are applied randomly to the beginning and the ending of the reads as well. The three datasets and ground-truth transcripts are publicly available and can be downloaded from http://doi.org/10.5281/zenodo.4019965.

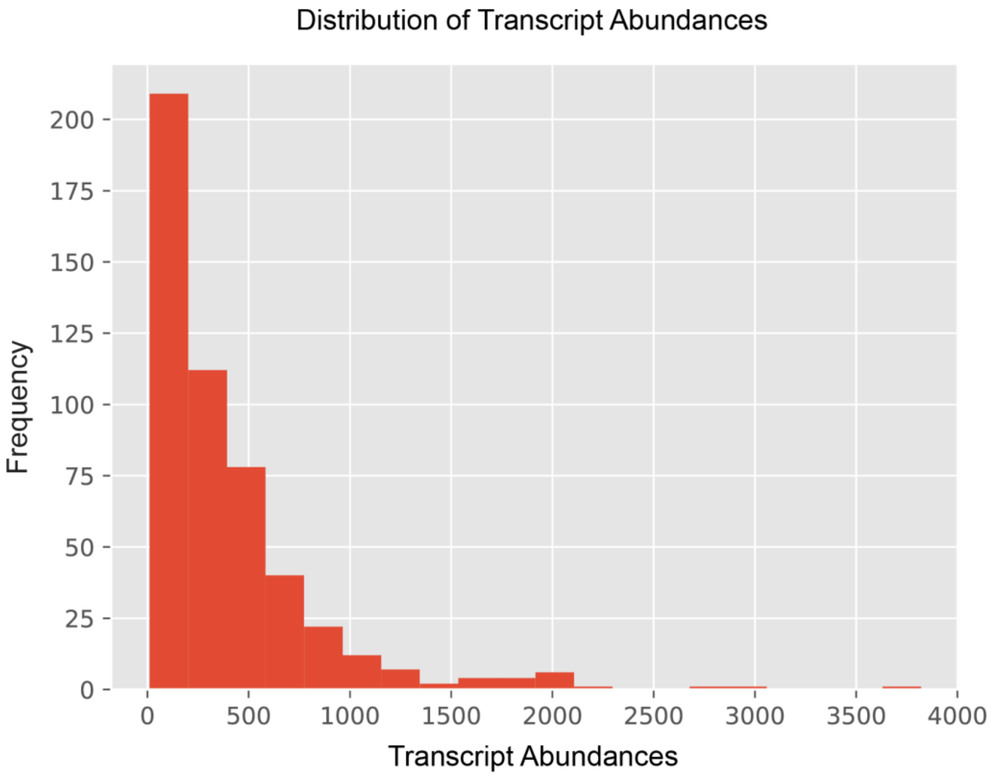


**Supplementary Figure 3:** Distribution of the Transcript Abundances for the 200K Simulated Dataset.

**SIRV Datasets:** The Lexogen SIRV dataset (https://www.lexogen.com/sirvs) is a synthetic RNA spike-in control and consists of four sequencing runs. Four libraries were constructed from the Lexogen SIRV E0 (two technical replicates), E1, and E2, respectively. Each library was sequenced on the PacBio RS II platform.

**FMR1 Dataset:** The FMR1 Iso-Seq dataset (http://doi.org/10.5281/zenodo.185011) consists of amplicon FMR1 cDNA from three premutation and three control brain samples. Each individual was prepped independently and sequenced on three SMRT Cells each on the PacBio RS II platform. The methodology of generating FMR1 dataset is fully described in Tseng et al. (2017) paper[24].

**Liver Dataset:** Liver dataset (https://www.pacb.com/blog/data-release-whole-human-transcriptome) consists of full-length transcriptome of human liver generated by Iso-Seq sample preparation protocol.

**Sequel II Dataset:** The Universal Human Reference (UHR) Iso-Seq dataset consisted of two SMRT Cell 8M runs. The cDNA library was a whole transcriptome Iso-Seq library generated using the Clontech SMARTer cDNA kit followed by SMRT bell library preparation. Data is publicly available and downloaded from https://downloads.pacbcloud.com/public/dataset/UHR_IsoSeq/.

**Supplementary Note 6: I-CONVEX Pipeline**

To run I-CONVEX on a given genome reads dataset, first move the fasta file to the Clustering folder and rename it to ***reads.fasta***. Then execute the following commands in order to obtain **pre-clusters**:

**Pre-clustering**

```
ClusteringReads/Clustering$ python SplitFile.py
ClusteringReads/Clustering$ chmod 777 commands.sh
ClusteringReads/Clustering$ ./commands.sh
```

The result is a Cluster folder containing subfolders, each of which represents a pre-cluster. Now, the following commands run the *clustering via convexification* module on pre-clusters:

**Clustering via Convexification**

```
ClusteringReads/Clustering$ python CreateConvexScript.py 8
ClusteringReads/Clustering$ ./run_convex.sh
ClusteringReads/Clustering$ python CollectTranscripts.py
```

The last command stores the final collected transcripts on the ***final_transcripts.fasta*** file.

To run all parts of the algorithm step by step, or change the hyper-parameters such as $k$, $d$, and $b$, follow the advanced version instructions on https://github.com/sinaBaharlouei/I-CONVEX.
